# Affordable and Effective Optokinetic Response Methods to Assess Visual Acuity and Contrast Sensitivity in Larval to Juvenile Zebrafish

**DOI:** 10.1101/2021.04.26.441419

**Authors:** Alicia Gómez Sánchez, Yolanda Álvarez, Basilio Colligris, Breandán N. Kennedy

## Abstract

**Background:** The optokinetic response (OKR) is an effective behavioural assay to investigate functional vision in zebrafish. The rapid and widespread use of gene editing, drug screening and environmental modulation technologies have resulted in a broader need for visual neuroscience researchers to access affordable and more sensitive OKR, contrast sensitivity (CS) and visual acuity (VA) assays. Here, we demonstrate how 2D- and 3D-printed, striped patterns or drums coupled with a motorised base and microscope provide a simple, cost-effective but efficient means to assay OKR, CS and VA in larval-juvenile zebrafish.

**Results:** In wild-type, 5 days post-fertilisation (dpf) zebrafish, the 2D or 3D drums printed with the standard OKR stimulus of 0.02 cycles per degree (cpd), 100% black-white contrast evoked equivalent responses of 24.2 or 21.8 saccades per minute, respectively. Furthermore, although the OKR number was significantly reduced compared to the 0.02 cpd drum (p<0.0001), the 2D and 3D drums evoked respectively equivalent responses with the 0.06 and 0.2 cpd drums. Notably, standard OKR responses varied with time of day; peak responses of 29.8 saccades per minute occurred in the early afternoon with significantly reduced responses occurring in the early morning or late afternoon, (18.5 and 18.4 saccades per minute, respectively). A customised series of 2D printed drums enabled analysis of visual acuity and contrast sensitivity in 5-21 dpf zebrafish. The saccadic frequency in visual acuity and contrast sensitivity assays, was inversely proportional to age, spatial frequency and contrast of the stimulus.

**Conclusions:** OKR, VA and CS of zebrafish larvae can be efficiently measured using 2D- or 3D-printed striped drums. For data consistency the luminance of the OKR light source, the time of day when the analysis performed, and the order of presentation of VA and CS drums must be considered. These simple methods allow effective and more sensitive analysis of functional vision in zebrafish.

## Background

The ability of researchers to effectively assess functional vision is critical to understanding the ontogeny of vision, the genetic and environmental mechanisms underlying impaired vison and the efficacy of therapeutic interventions (1, 2). The optokinetic response (OKR), or optokinetic nystagmus (OKN), is an innate behavioural response in humans, (3) primates (4), mammals (5) and teleosts (6). In clinical practice, the OKN is an objective measure of visual acuity, and can be evoked by presenting moving stimuli in front of patients by changing direction or size (7, 8). In natural environments, the OKR is essential for animals to hunt, feed and avoid predators. The OKR presents as a saccadic eye movement consisting of two phases: *i)* a slow eye movement following the stimulus, in the same direction as the stimulus; and *ii)* a rapid eye movement in the opposite direction to fixate on a subsequent stimulus. These movements help to stabilise the moving image presented to the retina. (9).

Here, we sought to generate simple and affordable tools for OKR assays in zebrafish and validate their efficacy in quantifying visual acuity and contrast sensitivity in larval and juvenile zebrafish. Zebrafish are widely used to investigate the biology of vision and blindness (10). Large clutches of embryos are readily obtained, which morphologically develop eyes within 24 hours, and by 5 days post-fertilisation (dpf) exhibit functional vision, including a robust OKR (11, 12). Commonly, OKR analyses in zebrafish only utilise one standard stimulus *i.e.* a drum of 0.02 cpd (*e.g.* 1 cm width stripes) and 100% black and white contrast stripes (13, 14). This is not sufficient to detect subtle impairments in vision. One approach to more thoroughly vision evaluations, is to vary the optokinetic stimulation. By varying the width of the stripes, visual acuity is measured efficiently (15). Altering the extent of contrast between stripes enables the measurement of contrast sensitivity (16). Such assays have previously been successfully performed in zebrafish by specialists often using automatic or semi-automated OKR stimulators and specialised software (15–17). However, such bespoke equipment is often inaccessible or unaffordable to many research groups. Here, we describe a simple and affordable method to assess visual acuity and contrast sensitivity in zebrafish using 2D and 3D printed striped patterns/drums to quantify OKR, VA and CS in larval and juvenile zebrafish.

## Results

### 2D and 3D Printed Visual Acuity Patterns Elicit Equivalent OKR Responses in 5 dpf Zebrafish

The manual OKR equipment set-up (***Fig. 1***) permits simple exchange of stimulus patterns to measure visual acuity. This apparatus was assembled using a microscope (***Fig. 1A***) to observe zebrafish eye movements, a light source (***Fig. 1B***) and an electronic motor connected to a 6 cm rotating circular base (***Fig. 1C***). 2D or 3D printed stimulus drums (***Fig. 1D***) were placed on the circular base which was rotated electronically to evoke eye movements. A standard OKR pattern of 0.02 cpd, (100% contrast) (***Fig. 1E***) and customised 0.06 and 0.2 cpd patterned stimuli (***Fig. 1D***) were produced by 2D or 3D printing (*see Methods and Additional data 1 for full details on OKR assembly)*.

**Fig 1.**
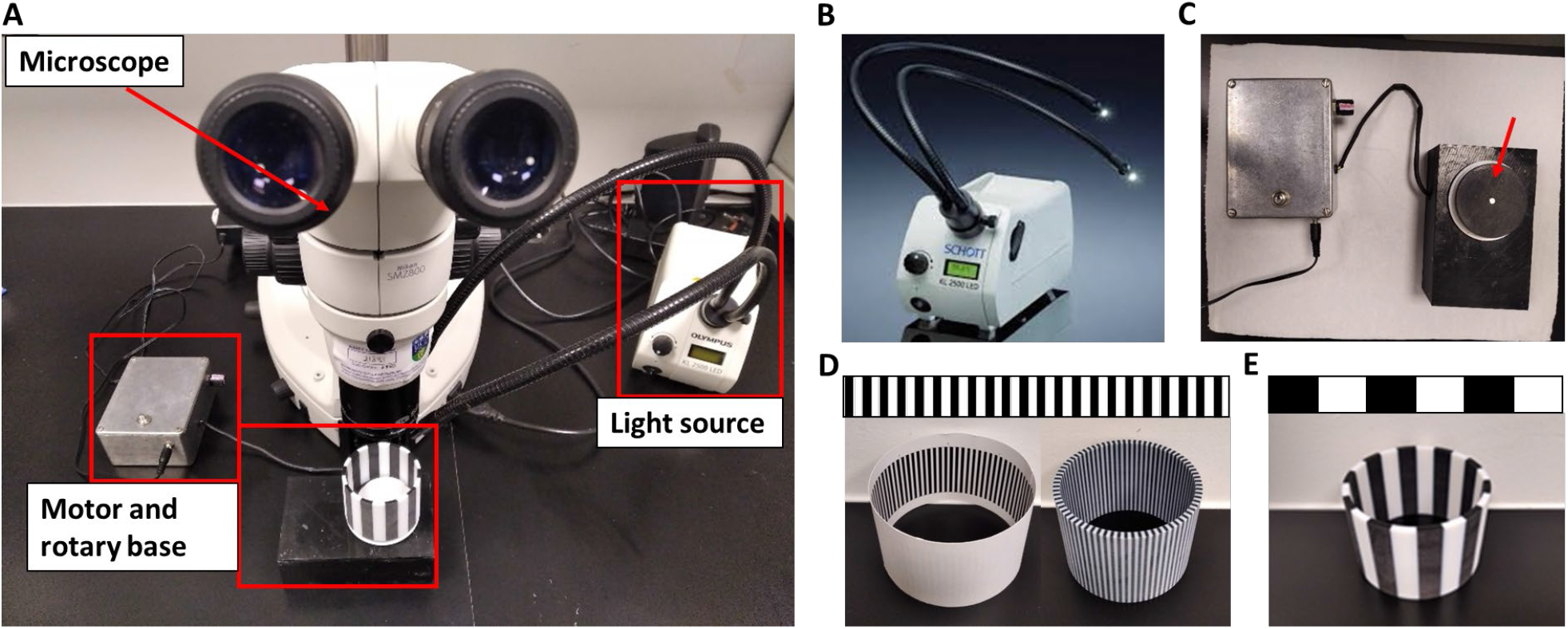
Optokinetic Response Equipment. A. Fully assembled Optokinetic Response apparatus. B. KL 2500 LED Schott light source C. Motorised rotary base to assemble the OKR drums (arrow). D. Optokinetic Response 2D (left) and 3D (right) drums of 0.2 cpd. E. Optokinetic Response 3D drum of 0.02 (standard OKR).

Visual acuity analysis with 2D and 3D printed drums was performed on 5 dpf zebrafish larvae (∼123.5 hours post-fertilisation - hpf) using *Protocol I* (see Methods) (***Fig. 2***). The OKR responses evoked by the 3D and 2D-printed drums were equivalent. More specifically, the OKR activity with the 0.02 cpd 2D-printed pattern (24.2 saccades per minute) was equivalent to the 21.8 saccades per minute evoked with 0.02 cpd 3D-printed drum (***Fig 2***). Similarly, there was no significant difference between the 7.5 and 5.3 saccades per minute, respectively produced by the 0.06 cpd 2D and 3D-printed drum (***Fig 2***). At the highest spatial frequency tested, 0.2 cpd, the number of saccades evoked by the 2D (7.9 saccades per minute) and 3D- (5.8 saccades per minute) drums also showed no significant difference. Therefore, both 2D and 3D printed drums can be used to measure the visual acuity of 5 dpf zebrafish larvae, the 3D-printed drums offering a more durable, but more costly option.

**Fig. 2.**
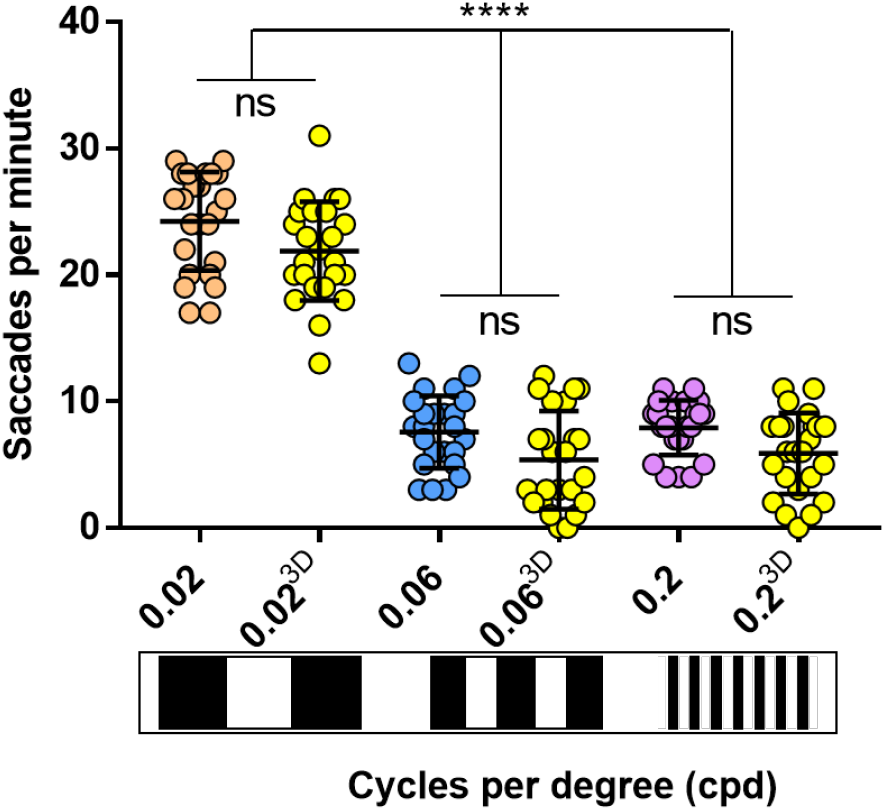
2D drums evoke same saccades frequency as 3D drums. Visual acuity responses obtained with 3D drums (yellow dots) don’t vary with respect of cardboard-printed drums (0,02 cpd, orange dots; 0,06 cpd, blue dots; 0,2 cpd, purple dots). 0.06 and 0.2 cpd responses evoked with 2D and 3D-printed drums were significant lower than standard OKR activity (0.02 cpd) with 2D and 3D-printed drums. Data were analyzed by one way ANOVA and Bonferroni’s multiple comparison test, where ns is no significative difference (p>0.05) and ****=p<0.0001. Error bars indicate standard deviation. 3 replicates of 8 larvae per each drum, n=24.

### The Zebrafish Larval OKR Response is Modulated by Time of Day and Luminance Levels

To determine if the zebrafish larval OKR has diurnal variations, the number of saccades generated with the standard 3D-printed OKR drum (0.02 cpd) was determined at 7 timepoints distributed throughout the light phases of the standard 14-hour light: 10-hour dark cycle (***Fig. 3A***). At 5 dpf, the trend observed was an increasing number of saccades until the afternoon with a subsequent drop in response (***Fig 3A***). The highest OKR response (29.8 saccades per minute) was observed at *early afternoon*/127.5 hpf, which was significantly higher (p=0.0001) than the OKR responses observed at *early morning*/121.5 hpf (18.5 saccades per minute) or at *late afternoon*/129.5 hpf (18.4 saccades per minute). The *midday* and *early afternoon* responses on 5 dpf (125.5 and 127.5 hpf, respectively) were significantly greater than the corresponding time of day responses at 4 dpf (100.5 and 103.5 hpf, respectively).

**Fig 3.**
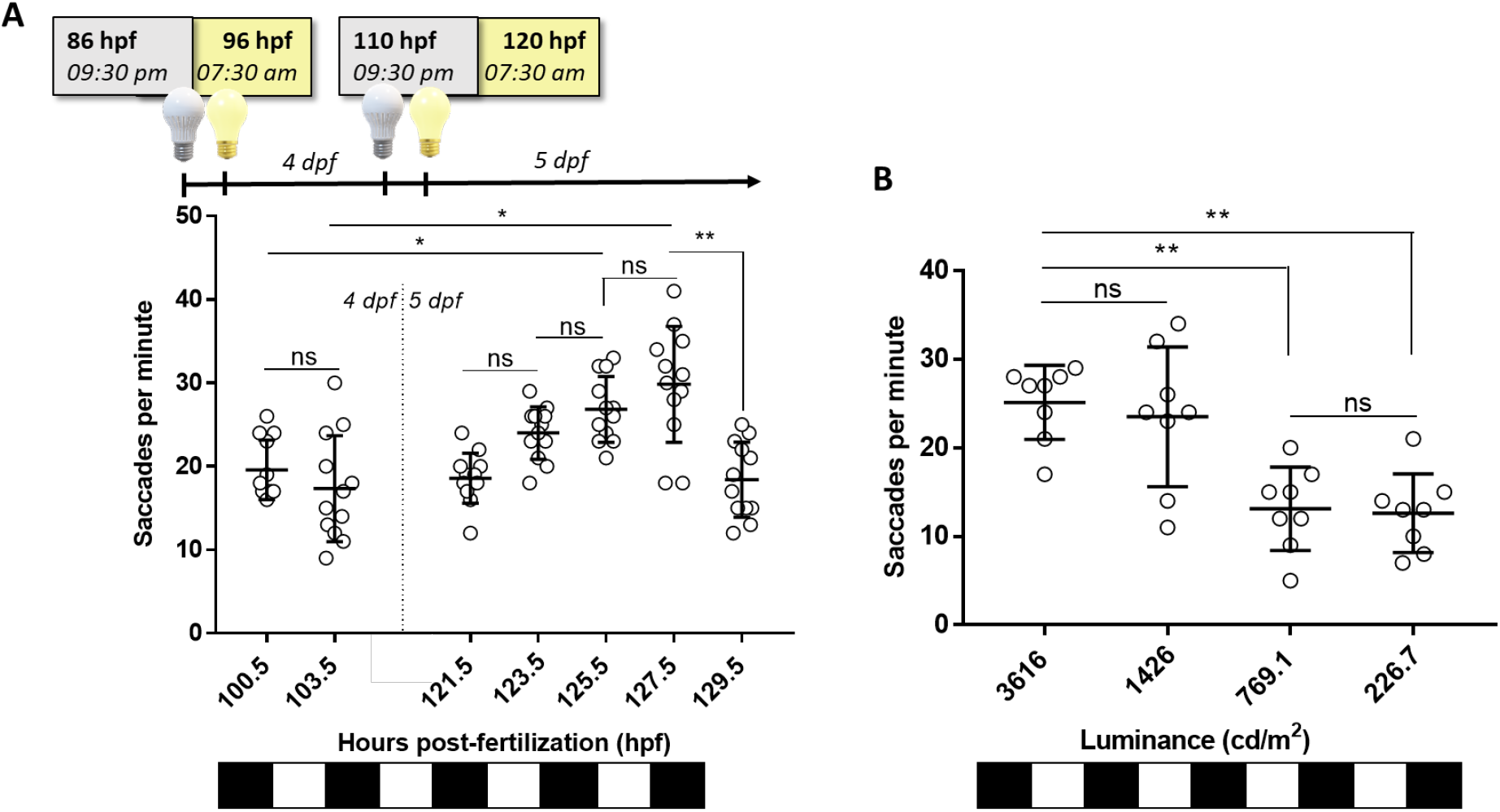
OKR Response is Modulated by Time of Day and Luminance Levels. A. Standard Optokinetic Response (0.02 cpd, 100% black contrast, 3616 cd/m2) at different timepoints along 4 and 5 dpf. Equivalent times (100.5 vs. 125.5 hpf; 103 vs. 127.5 hpf) show an increase OKR between 4 and 5 dpf. Higuest Optokinetic Response yields at 127.5 hpf. B. Standard Optokinetic Response (0.02 cpd, 100% black contrast) at different levels of luminance at 125 hpf. Higher levels of luminance evoked a better response on zebrafish but at 1426 cd/m2, OKR response is more variable (SD=7.8 saccades per minute) than 3616 cd/m2 (SD=4.1 saccades per minute). Data were analyzed by one-way ANOVA and Bonferroni’s multiple comparison tests, where ns is no significative difference, (p>0.05), **=p<0.01 and ****=p<0.0001. Error bars indicate standard deviation. 1 replicate of 12 larvae per each timepoint, n=12.

To evaluate if the 5 dpf OKR behaviour varied with brightness intensities, the standard OKR was assessed under luminance ranging from 226.7-3616 candelas per square meter (cd/m^2^) (***Fig. 3B***). The largest OKR activity occurred at 3616 and 1426 cd/m^2^(25.1 and 23.5 saccades per minute, respectively). The responses at 769.1 and 226.7 cd/m^2^(13.1 and 12.6 saccades per minute, respectively) were significantly lower (p=0.0081 and p=0.0035, respectively) than at 3616 cd/m^2^. In summary, the larval OKR shows response variations based on time of day recorded and light intensity used.

### The 2D/3D-printed Striped Patterns Enable Discrimination of Visual Acuity and Contrast Sensitivity in Larval Zebrafish

Establishment of affordable visual acuity and contrast sensitivity assays offers researchers the potential to identify more subtle defects in zebrafish vision than using standard OKR drums. Thus, bespoke 2D-printed striped patterns of 0.04 and 0.1 cpd for visual acuity were generated (*see Methods for details*) and tested (***Fig 4***). At 123 hpf, using *Protocol I (see Methods)*, an increased number of stripes reduced the number of saccades per minute, but robust and reproducible responses were observed at each cpd tested (***Fig. 4A***). At 0.04 cpd, the OKR activity (15.3 saccades/minute) was significantly (p<0.0001) lower compared to the standard OKR pattern of 0.02 cpd (24.2 saccades per minute), but significantly higher than the response with the 0.06 cpd pattern (7.5 saccades per minute, p<0.0001). The average saccades per minute with the 0.06 cpd pattern (7.6 saccades per minute) is similar to the 0.1 and 0.2 cpd pattern (6.9 and 7.9 saccades per minute respectively).

**Fig. 4.**
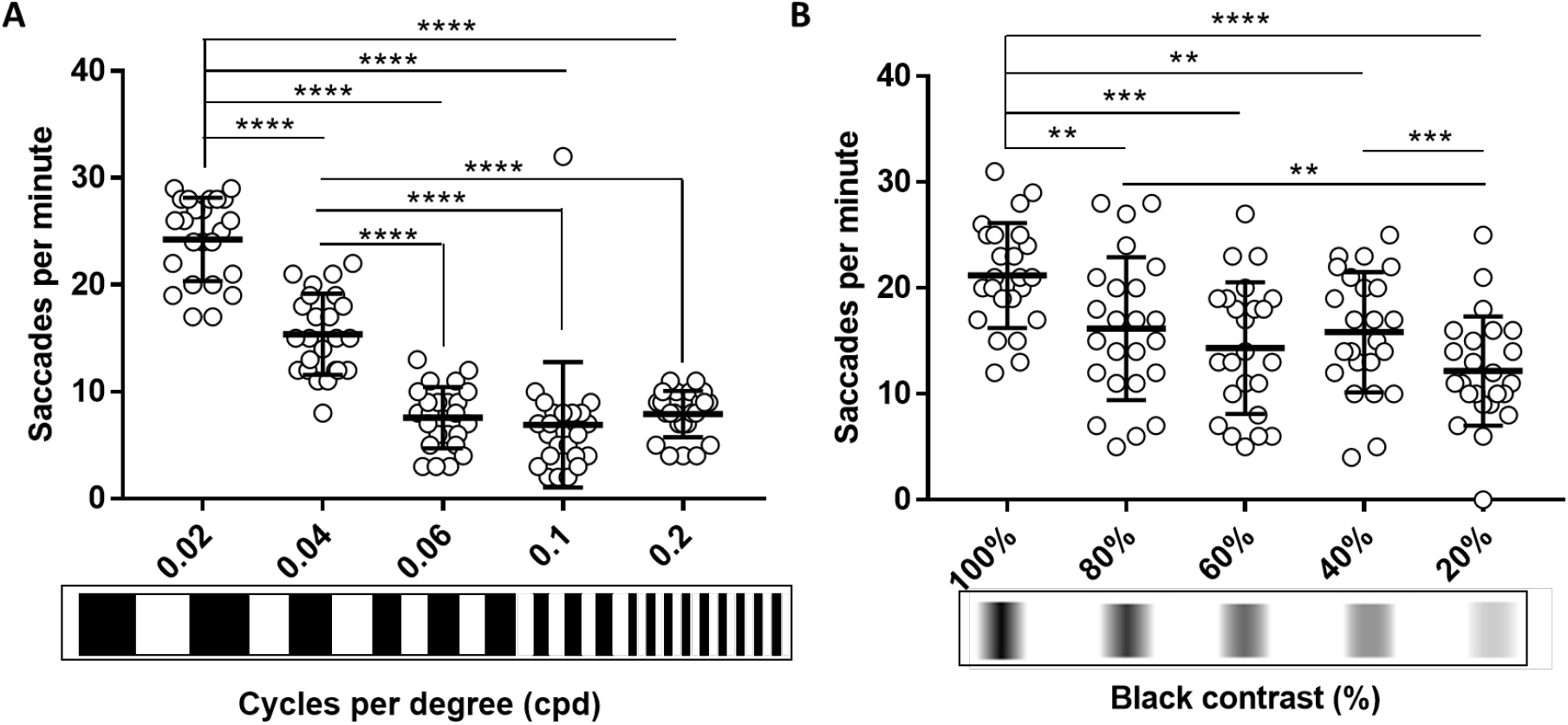
The 2D-printed Drums Enable Discrimination of Visual Acuity and Contrast Sensitivity in Larval Zebrafish. A. 5 dpf wild-type zebrafish larvae Visual Acuity decrease progressively when width of the stripes is reduced compared to standard OKR and 0.04 cpd. From 0.06 cpd, response is constant. B. 5 dpf wild-type zebrafish larvae Contrast Sensitivity decrease slowly when contrast between black-white stripes is lowered compared to standard OKR. 20% black contrast evokes the lowest response. Data were analyzed by RM one-way ANOVA and Bonferroni’s multiple comparison test, where ns is no significative difference, (p>0.05), **=p<0.01, ***=p<0.001 and ****=p<0.0001. Error bars indicate standard deviation. 3 replicates of 8 larvae, n=24 larvae per each pattern.

Contrast sensitivity assays were also performed using 2D printed drums and *Protocol I* at 125 hpf (***Fig 4B)***. The OKR activity evoked by the 0.02 cpd patterns with decreasing contrast was significantly reduced (80%, p=0.0022; 60%, p=0.0001; 40%, p=0.004, and 20%, p<0.0001) compared to the standard OKR drum of 0.02 cpd and 100% contrast. For example, at 80% black-white contrast, the 16.1 saccades per minute were significantly lower (p=0.0022) than the 21.2 saccades per minute evoked with the standard OKR drum pattern (0.02 cpd). There were no significant differences in response between the 80% contrast pattern and the 60% or 40% contrast pattern. The response from the 20% contrast pattern was significantly lower than with the 80% and 40% contrast pattern (p=0.0091 and p=0.0003, respectively. In summary, the 2D-printed patterns provide a simple and affordable method to assess contrast sensitivity and visual acuity assays in zebrafish larvae.

### The Zebrafish Visual Acuity Response Shows Age-Dependent Variations

Using the 2D-printed patterns, we determined if the OKR-based visual acuity response varies with age in larval to juvenile zebrafish aged 6, 9, 12, 16 or 21 dpf. Interestingly, with *Protocol I* the measured VA responses decreased with age (***Fig. 5A***) for all tested patterns. The largest OKR response of 24.6 saccades per minute was achieved at 5 dpf with a pattern of 0.02 cpd frequency (***Fig. 5A***). The lowest OKR response, with absence of any saccadic eye movements (0 saccades per minute), was obtained with 16 and 21 dpf zebrafish using patterns of 0.2 cpd (***Fig. 5A***). With patterns of 0.02 cpd, the OKR was significantly reduced at 16 dpf (p=0.0016) and 21 dpf (p<0.0001) compared to 5 dpf larvae, with 62% and 76% reductions in eye saccades, respectively. With patterns of 0.06 cpd, the highest responses were observed at 5 and 6 dpf (6.2 and 6.5 saccades per minute, respectively), which significantly declined at 16 dpf (0 saccades per minute, p<0.0001) and 21 dpf (0 saccades per minute, p<0.0001) compared to 5 dpf. For the highest VA patterns tested (0.2 cpd, with highest number of stripes), the largest OKR response was observed in 5 dpf larvae (7.5 saccades per minute) and significantly reduced responses were observed in 9 (1.7 saccades per minute, p=0.0004), 16 (0 saccades per minute, p<0.0001) and 21 (0 saccades per minute, p<0.0001) dpf zebrafish. Note, that at 16 dpf, when responses to VA and CS drums dropped, fish immobilisation in methylcellulose during drum stimulation was more difficult compared to earlier stages. In addition to observing an age-dependent reduction in OKR at each cpd frequency, we also observed that the level of response with the 0.06 and 0.2 cpd patterns were much lower than with the 0.02 cpd standard drum (***Fig. 5A***). In *Protocol I*, the data is generated based on first testing larvae at the lowest spatial frequency, and subsequent testing in the next higher spatial frequency drum. Therefore, to assess whether the reduction in OKR response with drums of higher spatial frequency was due to adaptation to previous OKR stimuli, we repeated the assays at 5 and 16 dpf, using *Protocol II (see Methods for details*) where each fish was tested with only one drum pattern (***Fig. 4B***). In 5 dpf zebrafish, there was no significant difference in OKR response using *Protocol I or II* for 0.2 cpd pattern (***Fig. 5B***). There was a significant increase (p=0.0017) in OKR response of 5 dpf larvae with *Protocol II* compared to *Protocol I* with the 0.06 cpd pattern. (***Fig. 5B***). However, the *Protocol II* response of 10.6 saccades per minute with the 0.06 cpd pattern was still significantly lower (p<0.0001) than the 24.6 saccades per minute observed under *Protocol I* with the 0.02 cpd standard drum (***Fig. 5B***). In 16 dpf zebrafish, a slight but significant increase (p=0.044) in OKR response was noticed when *Protocol II* is compared to *Protocol I* with the 0.06 cpd pattern. With the 0.2 cpd pattern and 16 dpf zebrafish there was no significant difference using *Protocol I* or *Protocol II*. At 16 dpf, the 0.06 cpd response obtained with *Protocol II* (2.1 saccades per minute) is significantly lower (p=0.0204) than the 0.02 cpd response (9.3 saccades per minute). In summary, all the above suggests that VA measurements drop after 12 dpf. Additionally, care needs to be taken regarding a consistent order of testing the VA drums to avoid experimental artifacts.

**Fig. 5.**
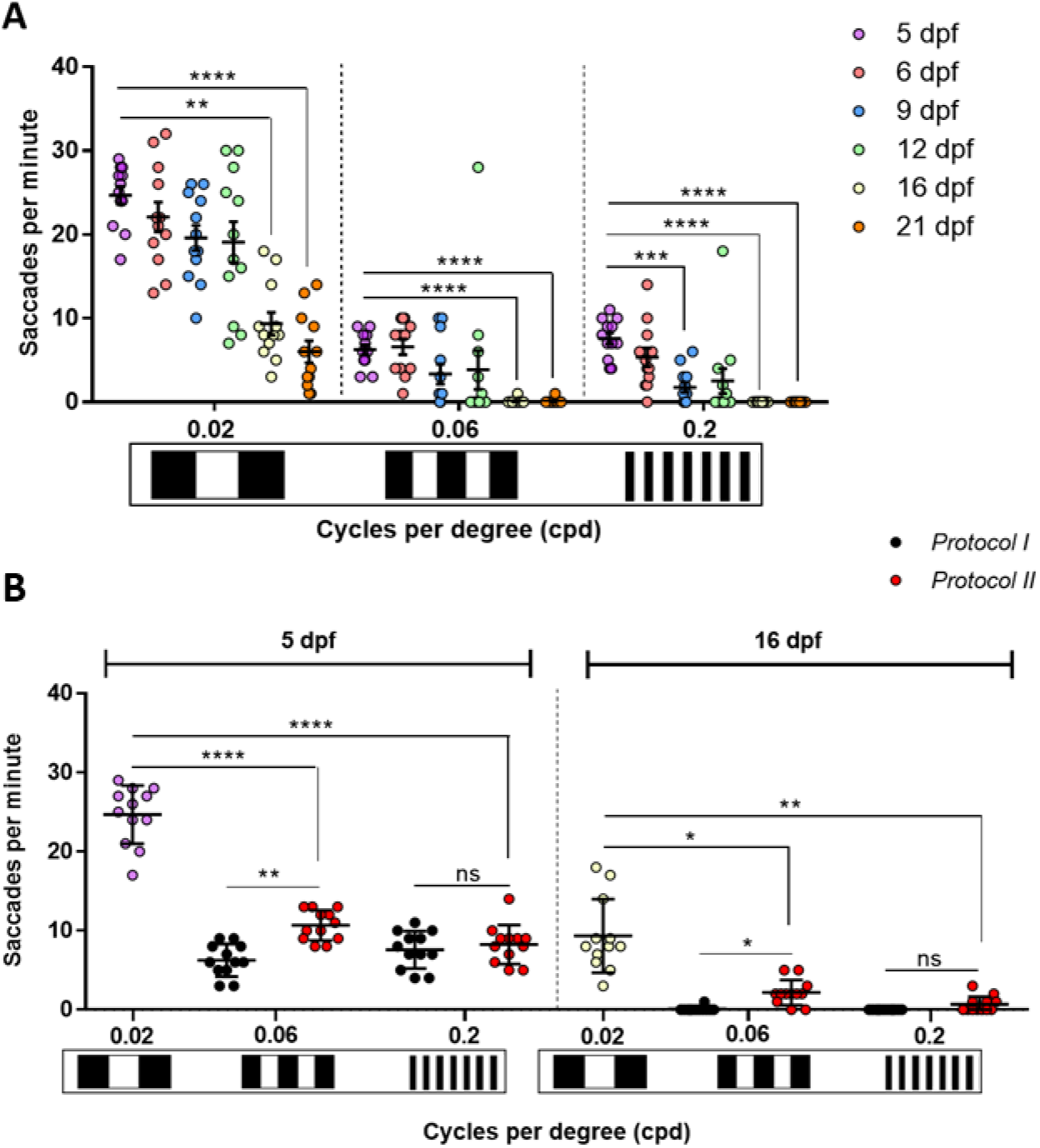
The Zebrafish Visual Acuity Responses Shows Age-Dependent Variations. A. Visual Acuity of zebrafish from 5 to 21 dpf drops significatively from 16 dpf following *Protocol I* at 0.02, 0.06 and 0.2 cpd. At 0.2 cpd, this decreased response is also remarkable on 9 dpf. 1 replicate of 12 larvae per each set of patterns. B. Comparison of Visual Acuities measured with *Protocol I* (black dots) and *Protocol II* (red dots) on 5 dpf and 16 dpf. 0.02 cpd responses at 5 dpf (purple dots) and 16 dpf (white dots) belong to *Protocol I* and *Protocol II* as it is the first pattern tested. There is no difference between both protocols except at 0.06 cpd where 5 dpf naïve larvae showed a higher number of saccades. Data were analyzed by RM one-way ANOVA and Bonferroni’s multiple comparison test, where ns is no significative difference, (p>0.05), **=p<0.01, ***=p<0.001 and ****=p<0.0001. Error bars indicate standard error of mean in A and standard deviation in B. 1 replicate of 12 independent larvae for each pattern, n=12.

### The Zebrafish Contrast Sensitivity Responses Show Age-Dependent Variations

Subsequently, we determined if the contrast sensitivity responses obtained using the 2D printed patterns displayed age-dependent variations. Using *Protocol I* and 0.02 cpd drums with 100% black-white contrast, the largest response of 25.2 saccades per minute was observed with 5 dpf larval zebrafish (***Fig. 6A***). Responses to these drums showed significant reduction with age, but reproducible visual behaviour responses were still observed with 16 and 21 dpf juvenile zebrafish (9.2 saccades per minute, p=0.0007; and 6 saccades per minute, p<0.0001, respectively). Similarly, with the 20% contrast drums, the largest responses were observed with 5 and 6 dpf (11.5 and 16 saccades per minute, respectively) larvae. Numbers declined with age and significant reductions were observed in 12, 16 and 21 dpf juveniles (3.7 saccades per minute, p=0.04; 1 saccadic per minute, p=0.0019; 0.1 saccades per minute, p<0.0001, respectively). As mentioned earlier, fish immobilisation and saccade counting in older fish is less consistent. Again, we utilised *Protocol II* to determine if reduced responses were due to desensitisation to consecutive stimuli. In 16 dpf zebrafish, there was no significant difference in OKR response using *Protocol I or II* when testing 20% contrast drums (***Fig. 6B***). In 5 dpf zebrafish, there was a significant increase (p=0.0002) in OKR response at 20% contrast when Protocol II is compared to *Protocol I* (***Fig. 6B***). Indeed, the *Protocol II* response of 24.3 saccades per minute with the 20% contrast drum is equivalent to the response observed under *Protocol I* with 100% contrast (***Fig. 6B***), suggesting the diminished CS response is due to desensitisation. In summary, our data suggests that CS responses decrease significantly after 9 dpf, and highlight the importance of strict consistency to be taken while testing different CS patterns on the fish to avoid confounding.

**Fig. 6.**
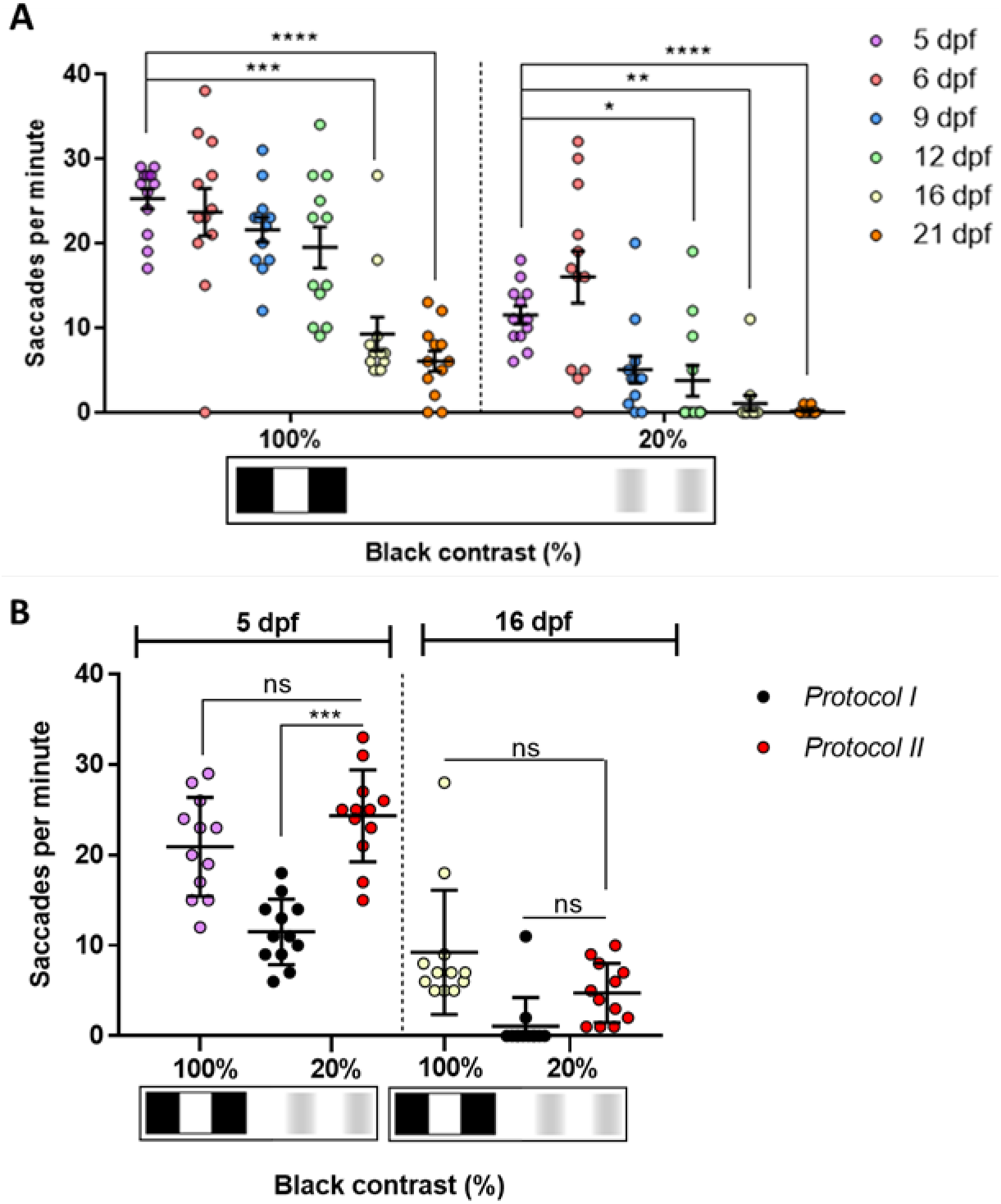
The Contrast Sensitivity response of juvenile zebrafish diminish with age. A. OKR response to 20% is significant at 12, 16 and 21 dpf. B. Comparison of contrast sensitivity responses measured with *Protocol I* (black dots) and *Protocol II* (red dots). 100% black contrast at 5 dpf (purple dots) and 16 dpf (white dots) belong to *Protocol I* and *Protocol II* as it is the first pattern tested. Responses of 20% black contrast with *Protocol II* are higher than when evoked with *Protocol I*. However, there are not differences between both protocols at 16 dpf. Data were analyzed by RM one-way ANOVA and Bonferroni’s multiple comparison test, where ns is no significative difference, (p>0.05), *p<0.05, **=p<0.01, ***=p<0.001 and ****=p<0.0001. Error bars indicate standard error of mean in A and standard deviation in B. 1 replicate of 12 independent larvae for each pattern, n=12.

## Discussion

### Affordability and Accessibility

The optokinetic response is a strong, innate visual behaviour that is very useful to characterise functional vision in zebrafish (12). We employed 2D and 3D printed patterns, of different stripe width or different black-white contrast, to effectively and affordably assay OKR, VA and CS in 5 to 21 dpf zebrafish. The remaining equipment required is accessible and affordable as suitable microscopes are commonly available in laboratories and the other components *e.g.* motor, light source and 2D/3D-printed patterns can be acquired easily and cost-effectively. Whilst automated or computerised devices were previously used to report optokinetic responses, those systems have high costs (up to €30,000), prohibitive to many research groups. Furthermore, computerised measurements of OKR, VA and CS, apply software to disaggregate the collective saccadic eye movement into eye velocity, gain or amplitude parameters (6, 15, 16). This requires establishment of thresholds based on algorithms and formulas using specialist programmes (6, 15). In summary, the manual OKR set-up described here, enables refined and accurate evaluation of OKR, VA and CS in zebrafish larvae, it is easy to use, does not require specialist software and is up to 10 times more affordable.

### Effectiveness and Sensitivity

With the 2D and 3D printed patterns, the magnitude of the 5 dpf VA response progressively decreased from the 0.02 cpd (standard OKR) to the 0.2 cpd (finest stripe width tested) pattern. This inverse relationship between saccadic response and stripe width agrees with previous studies using digitalised OKR set-ups and a 0.02 – 0.2 cpd range of visual acuity patterns (15, 16). Notably, those studies, which pre-stimulated the larvae with a 0.06 cpd pattern before testing, reported 0.16 cpd as the highest VA pattern to evoke an OKR in 5 dpf larvae *(15, 16)*. However, with our 2D and 3D printed drums, an even finer VA stimulus of 0.2 cpd elicits reproducible OKRs of 5.8-7.8 saccades per minute, providing enhanced ability to identify more subtle visual impairment phenotypes.

### Diurnal Variability

There is clear evidence of dynamic anatomical and behavioural development of zebrafish vision up to 5 dpf (9, 18–20). A previous analysis of diurnal variations in OKR at 5 dpf (21, 22) showed no difference in the number of saccades evoked during the day at 122 hpf (*early morning;* 27 saccades per minute) and 134 hpf (*early evening;* 25 saccades per minute), but dropping to 0 saccades per minute at 137 hpf (*night*) (21). Another study performed diurnal OKR analysis at different timepoints (22) wherein, at 125 hpf the OKR gain peaked and then decreased progressively at 129 and 133 hpf. Here, we investigate the OKR response between 4 and 5 dpf using even shorter time intervals. We found a cyclic modulation of OKR activity from a base of 19.5 saccades per minute at 100.5 hpf (*mid-day)* at 4 dpf, reaching a peak of 29.8 saccades per minute at 127.5 hpf (*early afternoon*) on 5 dpf and then troughing at 18.4 saccades per minute at 129.5 hpf (*late afternoon*) on 5 dpf. The peak responses at 125.5 and 127.5 hpf (26.8 and 29.5 saccades per minute) and diminished response at 129.5 hpf (18.4 saccades per minute) are consistent with Huang *et al* (22). This diurnal variation may be attributed to circadian rhythms that drive diurnal and nocturnal behaviours in zebrafish (22, 23). In summary, a more extensive characterization at shorter times post-fertilization demonstrates significant diurnal variations in the OKR and highlights the importance of carefully controlling the time of day when OKR analysis is performed.

### Light Variability

OKR gain, the ratio between eye velocity and stimulus velocity during the slow saccadic phase, was previously reported to increase with luminance from 0.38 cd/m^2^up to 388 cd/m^2^levels (16). Here, we demonstrate that higher luminance levels of 769.1 to 3616 cd/m^2^ increase the saccadic frequency from 13.1 to 25.1 saccades per minute. Notably, we did not, as in previous studies, measure the luminance from where the stimulus was projected (16). Instead our luminance was measured at the position of the fish in the methylcellulose to measure the ambient illumination surrounding the fish more accurately. In summary, the luminance of the light source must be measured and controlled during all analyses to avoid this confounding variable which affects saccadic frequency.

### Contrast Sensitivity and Visual Acuity Detection

Previous VA studies on 5 dpf larvae report that the magnitude of the OKR gain or eye velocity between 0.02 and 0.2 cpd was indirectly proportional to spatial frequency (15, 16). More specifically, an eye velocity of 4 degrees/second at 0.05 cpd reduced to 0 degrees/second (no eye movements) at 0.2 cpd (12). At the highest drum velocity tested (22.5 degrees per second) a gain response of 0.2 (max. gain=1) at 0.02 cpd decreased to 0.025 at 0.16 cpd, however, the gain peak of 0.3 was reported at a mid-frequency of 0.06 cpd (16). The VA responses with the 2D printed patterns concur with this spatial frequency-dependency as evidenced by 24.2 saccades per minute at 0.02 cpd reducing to 7.9 saccades per minute at 0.2 cpd, the latter response contrasting with no eye movements using the computerised OKR hardware. Thus, the OKR set-up described here emulates VA responses of automatic devices (15, 16) and furthermore, it detects quantifiable responses at higher spatial frequencies (12, 13). CS analysis using the 2D-printed drums (20-100%) at 5 dpf show a similar trend as previously reported with computerised set-ups (0.7 to 100% contrast). A higher number of OKR saccades or greater OKR gain is observed as the black-white contrast increases (16, 24). Notably, these computerised devices reported a low gain (16) and no eye movements (24) at 20% black-white contrast. However, our manual OKR set-up evokes reproducible OKR saccades of 12.1 per minute at the 20% black-white contrast. Hence, our affordable 2D-printed drums can elicit OKR responses that discriminate higher visual acuity frequencies and lower black-white contrast enabling more sensitive detection of VA and CS in 5 dpf zebrafish.

### Age Variability

OKR analysis in juvenile zebrafish older than 5 dpf was previously reported using computerised OKR set-ups (6, 12, 16). Orger *et al* (12) used a lower drum velocity (10 degrees per second) and described the standard OKR activity at 7 dpf, showing robust eye saccades through a motion detection OKR. Beck *et al* (6) investigating the OKR phases from 5 to 35 dpf, found that at 50 degrees per second drum velocity, gain decreased in all tested ages (5 to 35 dpf). Here, we use the 2D-printed drums to describe the saccadic frequency of 5 to 21 dpf zebrafish based on spatial frequency (***Fig. 5A***). As zebrafish became older, the saccadic frequency was decreased when spatial frequency was increased. We obtained quantifiable responses at all tested ages except at 16 and 21 dpf using our 0.06 and 0.2 cpd patterns. This reduction when zebrafish were older, was also observed using 100% and 20% black-white contrast 2D-printed patterns in 5 to 21 dpf zebrafish (***Fig. 6A***). At 6 dpf, Rinner *et al* (16) previously reported that OKR gain with 100% black-white contrast was approximately 0.7, decreasing to 0.3 with 20% black-white contrast. This response is similar to what we obtained at 6 dpf using the 2D-printed patterns, where 23.6 saccades per minute were obtained at 100% black-white contrast, decreasing to 11.5 saccades per minute at 20% black-white contrast. In summary, our data suggests that manual VA/CS analysis, using 2D-printed patterns, can be used to detect spatial frequency and contrast discrimination by zebrafish larvae at 6, 9, 12, 16 and 21 dpf.

### Protocol Variability/Desensitisation

The significant drop of VA and CS response observed after 16 dpf may be explained by the use of methylcellulose to immobilise the larvae, but which is also reported to hamper oxygen exchange in zebrafish older than 7 dpf and to decrease the OKR gain (9).

Studies on adult zebrafish, aged between 4-16 (25) and 12-24 (17) months, placed the fish further from the stimulus, *i.e.* 7.3 cm (25) and 19.5 (17) cm *versus* the 3 cm used here. According to the visual acuity concept and VA examinations in children (26), the eye to stimulus distance should be increased with age, which suggests that 16 dpf could be a *“key”* time-point to increase the distance stimulus-eye in zebrafish.

We considered that habituation could also account for reduced VA/CS at older stages. However, using 16 dpf naïve larvae (*Protocol II*), responses at highest spatial frequencies (0.2 cpd) and lowest contrast (20% black-white contrast) were similar as those tested in 16 dpf in *Protocol I*. Our overall interpretation is that, in general, *Protocol I* is more suitable to conduct VA/CS studies in zebrafish larvae due to ethical considerations to reduce the number of individuals used, while enabling follow-up of VA or CS multiple data obtained from a single specimen. *Protocol II* could be however more suitable, if obtaining maximum responses at 5 dpf is relevant for the study.

## Conclusions

The OKR set-up described here can be easily and cost-effectively acquired to measure OKR. Our 2D-printed patterns can reliably and feasibly quantify VA and CS response in zebrafish larvae from 5 to 16 dpf. The age of the fish used, the time of the day the assay performed, the light levels within the fish position and pre-stimulation can vary the OKR response and must be accurately determined for a consistent OKR, VA and CS analysis. The 2D/3D drums and methods described here can be utilised to identify and characterise more effectively zebrafish models with visual deficits.

## Material and Methods

### Zebrafish husbandry

Adult wild-type (*wt-Tübingen*) zebrafish were maintained in holding tanks on a 14:10 h light-dark cycle in a recirculating water system under environmental parameters averaging temperature of 28°C, conductivity of 1347 μS and pH of 7.1 (27). Adult *wt* zebrafish were fed shrimp and dry pellet food twice daily. After the noon feed, male and female adults were placed in breeding tanks and *wt* zebrafish embryos obtained by natural spawning, collected the next morning and raised in embryo medium (0.137 M NaCl, 5.4 mM KCl, 5.5 mM Na2HPO4, 0.44 mM KH2PO4, 1.3 mM CaCl2, 1.0 mM MgSO4 and 4.2 mM NaHCO3 with 1 ml methylene blue) until 5 days post-fertilisation (dpf). Larvae were fed: i) SDS 100 and paramecium from 5 to 10 dpf, ii) SDS 100, paramecium and shrimp from 11 to 20 dpf, and iii) SDS 200 and shrimp from 21 to 28 dpf. All experiments using animals were approved by ethical approval granted by the UCD Animal Research Ethics Committee (AREC).

### Optokinetic Response Equipment

A simple and affordable OKR apparatus (***Fig. 1***) was assembled with a Nikon SMZ800 microscope (Micron Optical) to observe zebrafish eye movements; an electronic motor (RS Radionics) connected to a non-patterned 6 cm rotating circular base in which the 2D printed striped stimulus pattern was placed (***Fig 1C***). A Schott KL2500 LED light source (Mason technologies) fitted with dual goose neck lightguides was positioned to illuminate inside the drum (***Fig. 1B***). The dimensions of the 2D-printed striped patterns, generated with MS PowerPoint® and printed on stock cardboard, were 3.4 cm high and 6 cm in diameter (***Fig. 1D*** and ***Additional file 1***). The visual acuity drums ranging from 0.02 - 0.2 cycles per degree (cpd) were chosen based on a previous publication (15). They were designed by changing the width of the 100% black and white contrast stripes to the calculated cycles per degree (cpd = *n° of cycles/360°)* when mounted on the rotating base. Contrast sensitivity drums ranging from 100 - 20% were generated by degrading horizontally from the lateral sides to the centre and then changing the transparency percentage of the centre of the black stripes with all retaining 0.02 cycles/degree. Additional visual acuity drums were printed with 3D printing technology in polylactic acid (PLA), a stronger thermoplastic (Materialise UK Ltd) and placed on rotating circular base (***Fig. 1E***). 3D drums were designed following the same parameters as 2D-printed patterns (height= 5 cm; diameter=6 cm; cpd=0.02, 0.06 and 0.2;>99% contrast).

### Luminance measurement

An LS-100 luminance meter (Konica Minolta) measured, in candela per square meter (cd/m^2^), the light reflected from the drum under different light intensity settings of the Schott 2500. The luminance meter was placed at 18 cm horizontally and 30 cm high from the centre of the Petri dish at 60° angle. We establish 4 measurements at 22.7, 12, 7 and 2%, corresponding to 3616, 1426, 769.1 and 226,7 cd/m^2^.

### Drum velocity

The drum base was rotated with a constant angular velocity of 100 degrees per second.

### Visual Acuity and Contrast Sensitivity Methods (Protocols I and II)

To measure saccades/minute, a 6 cm Petri dish with 9% methylcellulose (Sigma Aldrich, UK) diluted in embryo medium was placed inside the rotating drum. From another Petri dish, larval or juvenile zebrafish in embryo medium were randomly chosen and immobilised in the centre of the OKR Petri dish with 9% methylcellulose. Rotating the patterned drums 30 seconds clockwise, followed by 30 seconds counterclockwise at 100 degrees/second, evoked horizontal eye movements which were counted manually. The standard OKR utilised a drum with 0.02 cpd and 100% black-white striped contrast (***Fig. 3***, ***Additional file 1A***). For visual acuity assays, the OKR was performed with 2D printed patterns of 0.02, 0.04, 0.06, 0.1 and 0.2 cpd and 100% black stripe contrast for all cpd tested (***Fig. 4A***, ***Additional file 1A-E***). In the contrast sensitivity assays, OKR was performed with 2D printed patterns of 100%, 80% 60%, 40% and 20% black/grey-white striped contrast, and 0.02 cpd all percentages (***Fig. 4B***, ***Additional file 1F-I***). Drums were presented following that order, from lowest to highest spatial frequency and from highest to lowest black striped contrast. Two different protocols *(I and II)* were applied. In *Protocol I*, a zebrafish larva was randomly chosen, placed central of the 0.02 cpd drum and saccades per minute were counted. Subsequently and consecutively, the drum was replaced with one of higher spatial frequency (for visual acuity) or lower contrast (contrast sensitivity) and saccades per minute counted. After completing one set of drums, another larva was randomly selected and used to repeat the same drum sequence. In *Protocol II*, instead of presenting each drum of a series to the same larva, different specimens were used for each drum, i.e. larvae were naïve for OKR. In practice, a larva was analysed with 0.02 drum, saccades per minute were counted and next replaced by another larva which was subjected again to the same drum. This procedure was repeated for the rest of the drum patterns.

### Statistical analysis

Statistical analysis was completed using GraphPad Prism 7.00 software (GraphPad, San Diego, CA). One-way repeated measures ANOVA was employed to determine significant differences between groups followed by Bonferroni’s multiple comparisons test. Significance levels were set at p < 0.05.

## List of abbreviations

AREC: Animal Research Ethics Committee
Cd/m^2^: candela per square meter
Cpd: Cycles per degree
Cm: centimetres
CS: Contrast sensitivity
Dpf: Days post-fertilization
Hpf: hours post-fertilization
OKN: Optokinetic nystagmus
OKR: Optokinetic response
PLA: Polylactic acid
VA: Visual acuity
Wt: wild type

## Declarations

### Ethics approval and consent to participate

All experiments carried out on zebrafish were performed according to ethical approval granted by the UCD Animal Research Ethics Committee (AREC-Kennedy).

### Consent for publication

Not applicable

### Availability of data and materials

All data generated or analysed during this study are included in this published article (and its additional information files).

### Authors’ contributions

AGS carried out and analysed all the experimental data. YA supervised the study and provided contributions in drafting and writing the manuscript. BNK conceived, designed and supervised the study, and provided major contributions in drafting and writing the manuscript. All authors read and approved the final manuscript.

## Acknowledgements

The authors thanks to Dr. Rebecca Ward and Ailis Moran for their assistance and helpful discussions related to this project.

## Additional files

Additional file 1 (.pdf): 2D Visual Acuity and Contrast Sensitivity patterns. 2D-printed patterns to perform Visual Acuity and Contrast Sensitivity assays.

