## Additional file 1 for "Affordable and Effective Optokinetic Response Methods to Assess Visual Acuity and Contrast Sensitivity in Larval to Juvenile Zebrafish"

***Additional file 1.*** *2-D Visual Acuity (A-E) and Contrast Sensitivity (F-I) patterns*

Drums were cut by dotted line and assembled by the discontinuous line to build the drums.

cpd=cycles per degree; BC= black contrast

**A) 0,02 cpd 100% BC**

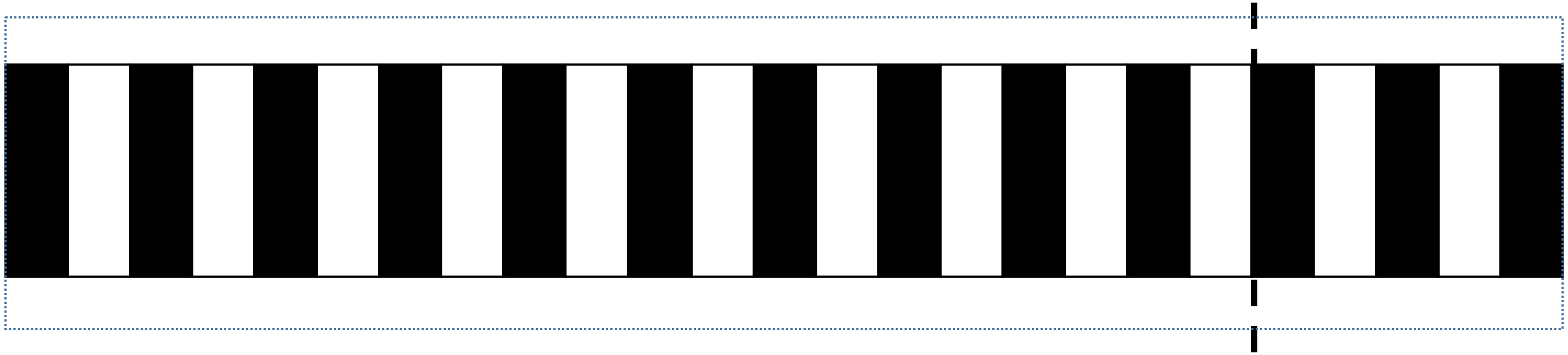

**B) 0,04 cpd 100% BC**

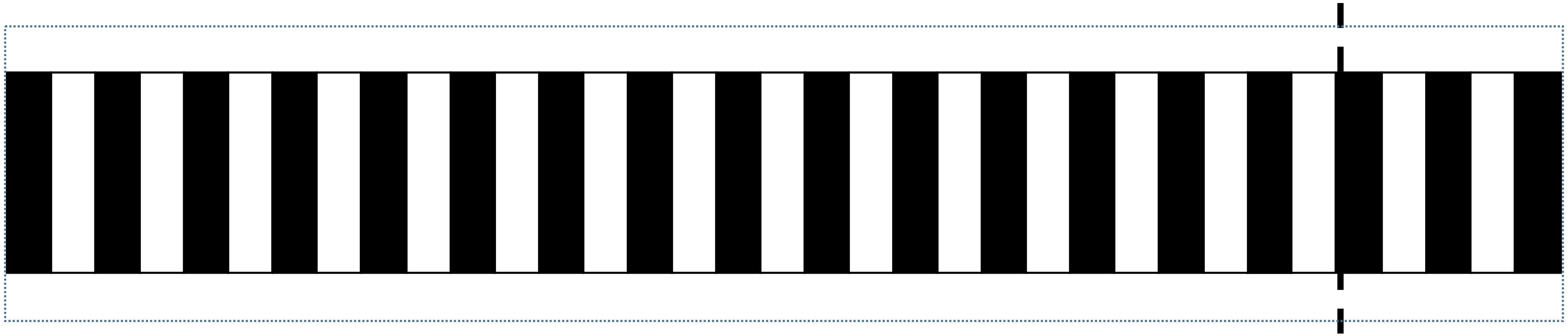

C) 0,06 cpd
100% BC

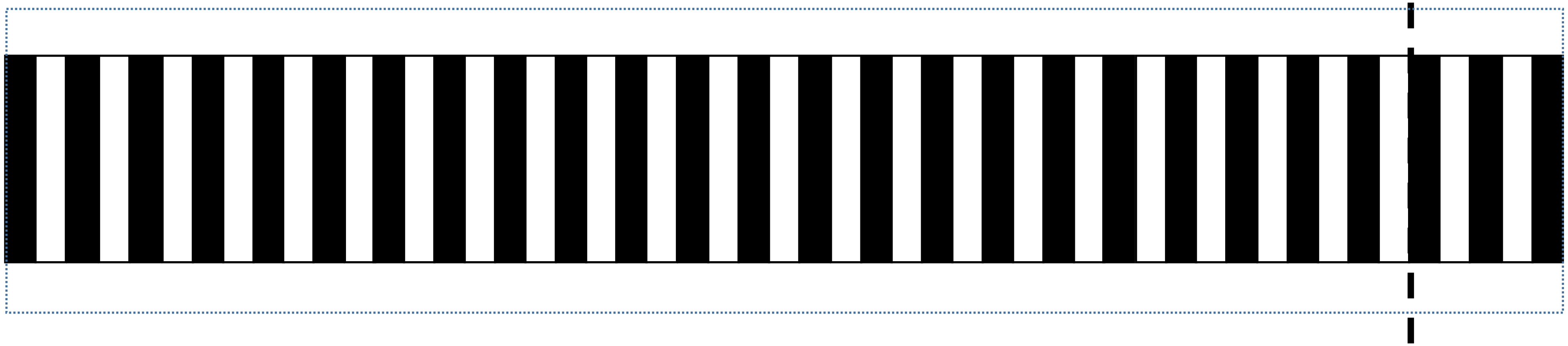

D) 0,1 cpd
100% BC

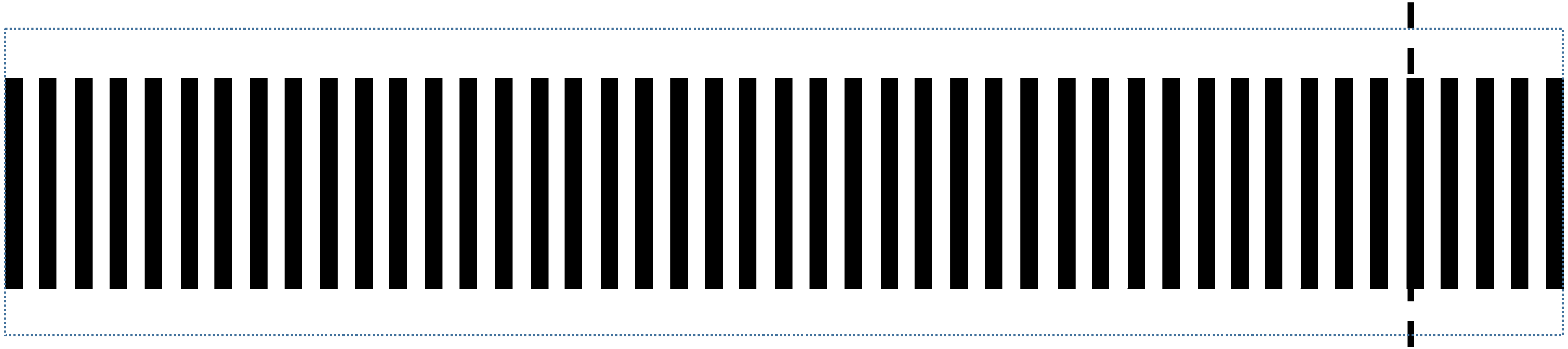

E) 0,2 cpd
100% BC

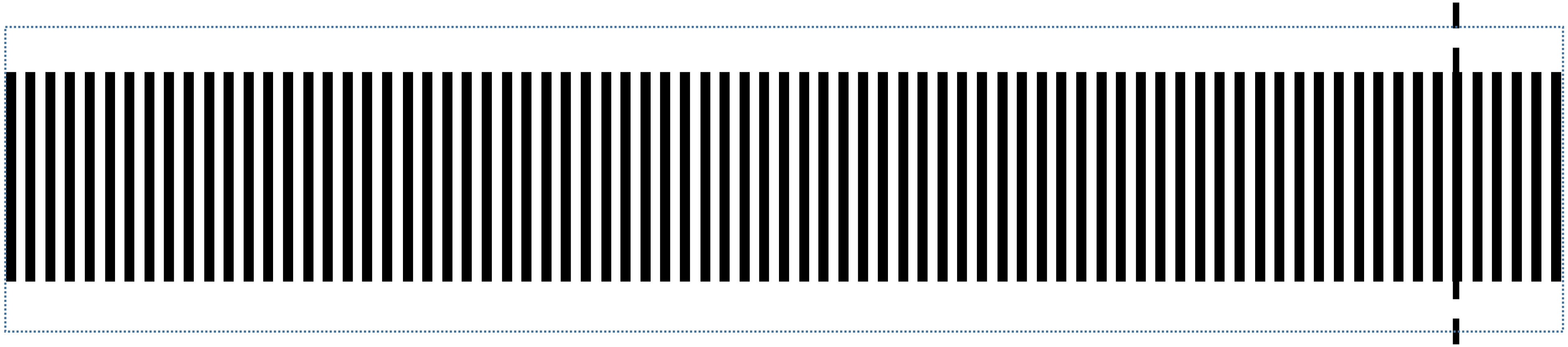

F) 80% BC
0,02 cpd

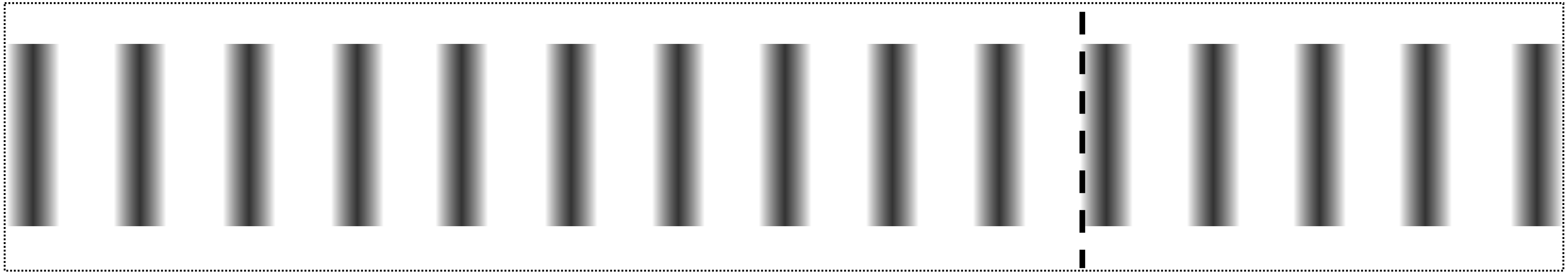

G) 60% BC
0,02 cpd

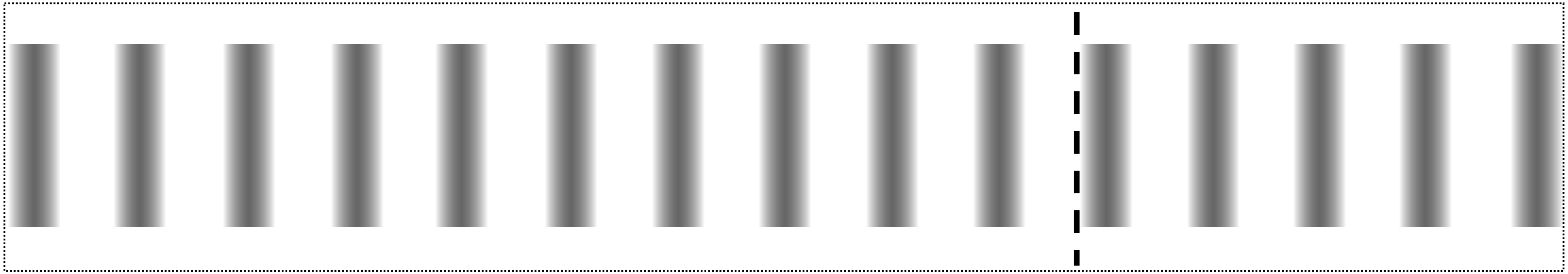

H) 40% BC
0,02 cpd

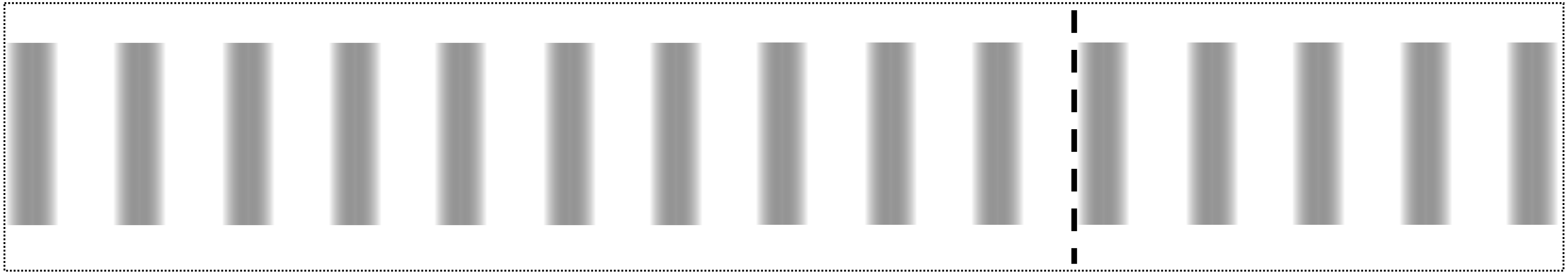

**l)** **20% BC**
**0,02 cpd**

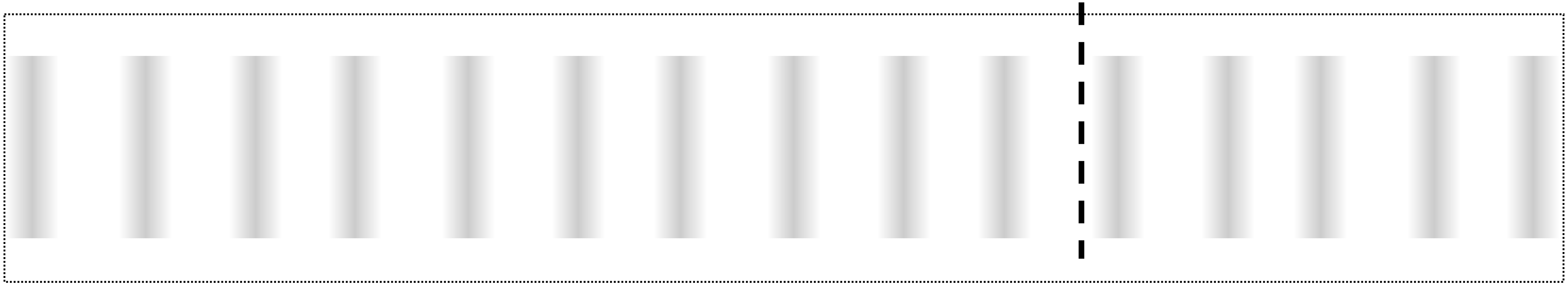
